# A Step-Emulsion Based Digital-RPA for Pathogenic Bacteria Detection

**DOI:** 10.1101/2024.04.07.588437

**Authors:** Jing jin, Ling Lu, Huicheng Chen, Yunhua Wang, Gouxia Zheng

## Abstract

Foodborne bacteria are major causes that affect human health. Development of new method that could achieve Rapid, sensitive and quantitative detection of pathogen is urgently needed. In this research, a step emulsion microfluidic,combined with droplet-based digital-RPA, was developed to detect *Vibrio parahaemolyticus*, a major seafood-borne pathogenic bacteria. Specific and rapid detection of *Vibrio parahaemolyticus* in 30 min has been achieved by this new device, with a detection limit of 10 CFU/μL, about 10-times lower than classical tube-based RPA. This device was demonstrated to be a promising tool for detection of pathogenic bacteria.

## 1. Introduction

Foodborne bacterial are major causes that affect human health, causing symptoms such as fever, abdominal pain, and vomiting. Most foodborne infections are caused by improperly handled and cooked food infected with pathogenic bacteria, including *Salmonella enterica* (*S. enteric*), *Escherichia coli* O157:H7 (*E. coli*), and *Vibrio parahaemolyticus* (VP).^1^ *Vibrio parahaemolyticus* is a curved rod-shaped halophilic bacteria. It is found worldwide, mainly in coastal or river mouth area. ^2^Consuming raw or undercooked seafood contaminated with *VP* can lead to acute gastroenteritis. Infection of VP is considered a major cause of seafood-borne diseases.^3,4^ Therefore, rapid, specific, reliable and quantitative examine methods are urgently needed to facilitate detecting and controlling VP in seafood, environmental or clinical samples.

Currently, regular techniques used in foodborne bacteria examine include bacterial culturing, biochemical testing, immunological testing, and real-time polymerase chain reaction (qPCR), among others. Bacterial culturing and biochemical testing are often used as a gold standard for detection. But they are time-consuming, labor-intensive, typically taking at least 3 days to obtain results, with relatively low resolution. Furthermore, these methods are hard to quantify the bacterial. qPCR technology is highly specific and sensitive. For example, the lowest detection limit for Vibrio parahaemolyticus can reach 10^4^ CFU/mL using qPCR.^5^ However, in qPCR quantitative analysis, samples are compared to a standard curve to determine targets levels, which resulted in a complicated process. Besides, expensive PCR equipment for temperature cycling and well-trained operator are need. Moreover, the detection process typically takes at least 2 hours. These factors hindered its application in point-of-care testing (POCT).^6^

Microfluidic technology integrates microchannels, microelectrodes, reaction chambers, microvalves, and microsensors on a single chip to manipulate, process, and analyze nanoliter or microliter volumes of liquid in microscale spaces, making it to be considered as a potential POCT technology platform.^7^ Droplet microfluidics, in particular, focuses on studying non-miscible multiphase flows that generate droplets within microchannels. It has been widely applied in single cell analyzing, molecular detection, drug delivery systems and other fields. ^8-10^As a potential POCT technology platform, the most prominent advantage of droplet microfluidic is that it does not require a standard curve for quantitative analysis. Microfluidic droplet technologies combined with nucleic acid amplification techniques have achieved low sample volume, high throughput, and high sensitivity in detection. ^11,12^ For example, SCHULER et al. designed the DropChip device, combining loop-mediated isothermal amplification (LAMP) to achieve absolute quantification of *E. coli*, with a detection range of 15-1500 DNA copies per microliter. ^13^NIE et al. utilized digital PCR combined with step emulsification microfluidic chips to detect the specific gene HER2 in breast cancer, achieving a sensitivity of 10 copies/μL.^14^

In this research, Recombinase Polymerase Amplification (RPA), an isothermal amplification technique, is utilized, which can amplify nucleic acids within 30 minutes at the optimal temperature range of 37°C to 42°C. RPA method offers many advantages of being rapid, sensitive, and suitable for use in POCT settings, making it a promising alternative to traditional diagnostic methods for foodborne pathogens. ^15^

Various techniques, including T-junctions, flow focusing, and step emulsification, have been developed to generate monodisperse droplets, enabling the production of highly monodisperse, high-throughput, and stable microdroplets. Among these methods, step emulsification is unique because it does not rely on precise flow rate control like others methods. The sizes of droplets generated by step emulsification are mainly determined by channel structure, making the manipulation easy and stable. Thus, it has incomparable advantages in generation droplets for POCT devices. ^16^

In this research, we developed a novel method for detection of VP. Step emulsification microfluidic chip combined with RPA is used to perform digital-RPA. In this method, DNA molecules in samples are divided into nanoliter-droplets according to the Poisson Distribution Law and then were amplified by RPA. By counting the number of positive and negative droplets, sample concentrations can be quantified absolutely without relying on standard curves. This approach allows for extremely sensitive detection of specific nucleic acid sequences within 25 minutes. This method increases the accuracy of target detection, providing a simple, low-cost, and powerful nucleic acid quantification technique. It offers a promising approach for quantitative detection of pathogen in resource-limited areas and POCT, without the need for sophisticated equipment, well-trained operator or complex procedures.

## 2 Experimental Methods

### 2.1 Bacterial DNA Extraction

All bacterial strains, including *Vibrio parahaemolyticus* (Vp), *Proteus* (Pro), *Vibrio aerogenes* (Va), *Escherichia coli* (*E*.*coli*), *Vibrio alginolyticus* (Vg), and *Pseudomonas aeruginosa* (Pa), were reserved in our lab. The strains were inoculate onto LB agar plates and cultured overnight. Single colony was picked and inoculated into liquid LB culturing medium and then cultured for 16 hours with constant shaking. Take 0.5 mL of bacterial culture for bacterial genomic DNA extraction using a DNA extraction kit. Simultaneously, bacterial concentration of *Vibrio parahaemolyticus* was counted by plate counting method.

### 2.2 Conventional RPA Detection of *Vibrio parahaemolyticus*

Conventional RPA detections were performed to amplify Vp-specific *tlh* gene using the TwistAmp exo kit, examining the specificity and sensitivity of probe and primers. The primers and probes are synthesized by Kingsray Biotech Company (Nanjing, China). The design sequences of primers and probes, as well as the reaction system, are shown in Table 1 and Table 2.

**Table 1.**
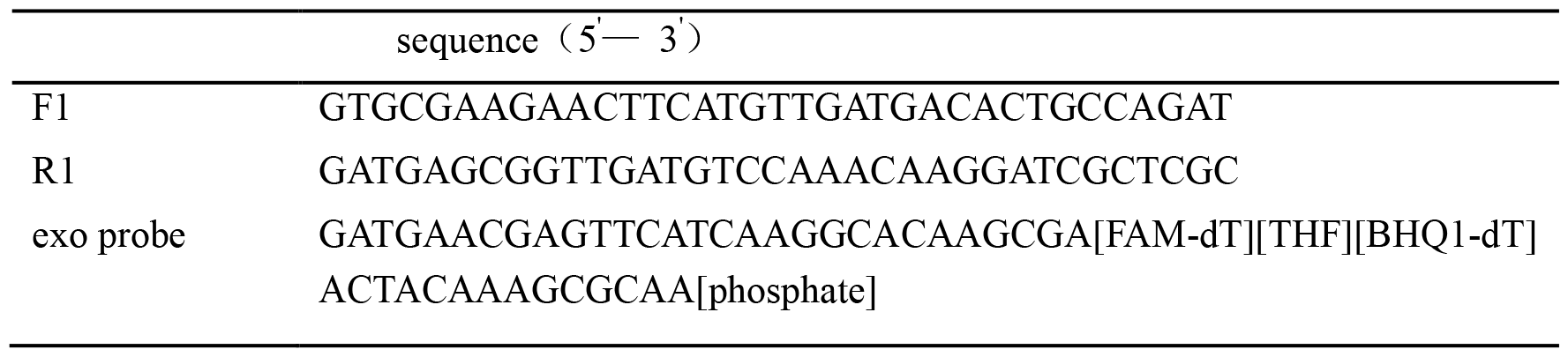
The primer sequences of tlh gene.

**Table 2.** The reaction system of RPA.

| Reagent | Volume |
| --- | --- |
| F3 (10μM) | 2.1μL |
| R3 (10μM) | 2.1μL |
| TwistAmp exo probe | 0.6μL |
| Primer Free Rehydration buffer | 29.5μL |
| DNA | 1.0μL |
| ddH<sub>2</sub>O | 12.2μL |
| MgOAc (280mM) | 2.5μL |
| Total volume | 50μL |

The amplification was performed according to TwistDx user manual. The probe concentration and reaction temperature for the traditional RPA system were optimized. Specificity and sensitivity were evaluated.

### 2.3 Design and Fabrication Step Emulsion Chip

The chip consists of droplet generation nozzle arrays, a microchamber to accommodate the droplets, an inlet and outlet (as shown in Figure 1). The heights of microchannels are 10-50 μm in different chipset and the height of micro-chamber is 200μm. The chip was made by classic soft lithography process as described elsewhere. Briefly, photoresist SU8-3035 (Microchem, USA) was spin coated onto a silicon wafer. The pre-designed patterns were transferred by photolithography process. Functional units that with different height were fabricated by double patterning. Polydimethylsiloxane (PDMS, Sylgard 184) monomer was mixed with the curing reagent at a ratio of 10:1 (w/ w) and was then degased for 1h in vacuum flask. The mixture was casted onto the SU-8 mold fabricated by soft lithography and then baked at 70°C for 45 min to get the PDMS sheets. The inlet and outlet hole were punched. The PDMS sheet was bonded to glass sheet after being treated with plasma cleaner for 1 min. The device was then baked at 60°C overnight to achieve irreversible sealing.

**Figure 1.**
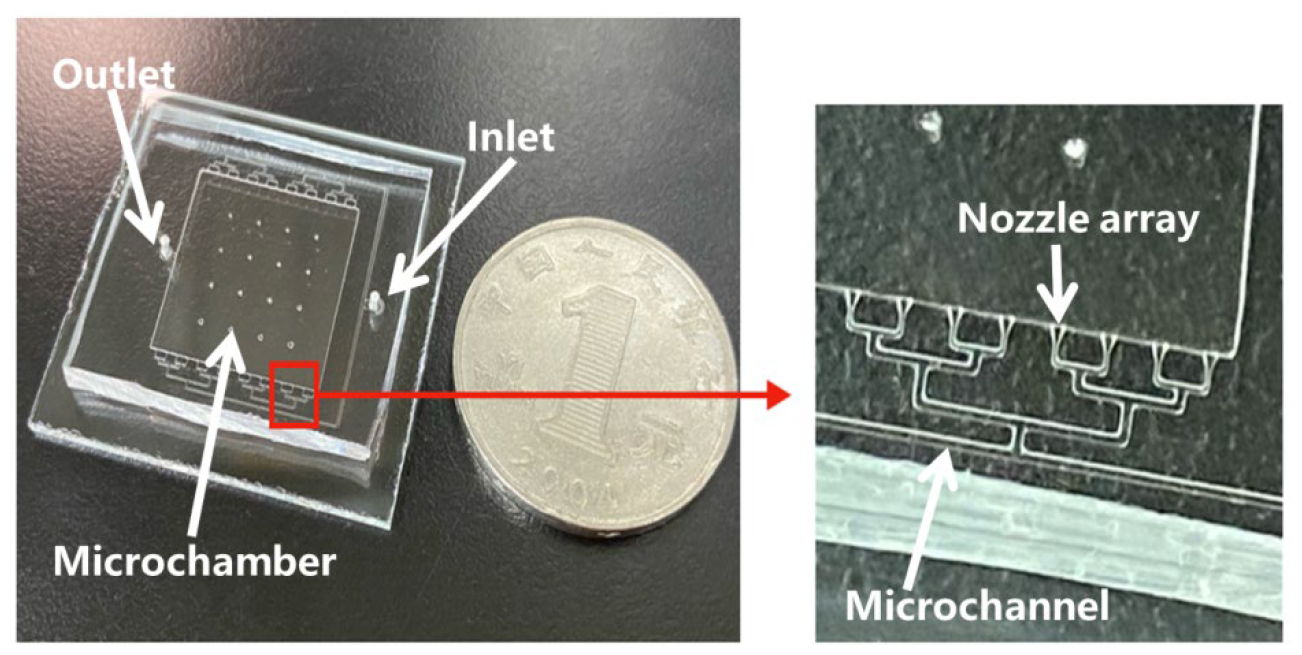
Step Emulsion Chip.

### 2.4 Characterization of the Chip

It was reported that the diameter of droplets generated in the step emulsification is related to the height of the steps at the nozzle. Different SE chips with channel heights of 10, 20, 30, 40, and 50 μm were designed to investigate the relationship between chip channel height and droplet diameter.In the step emulsification manipulation, the entire chip was initially filled with the continuous phase and then the dispersed phase is injected using a syringe to form monodisperse droplets. To investigate the impact of the flow rate of the dispersed phase on the droplet diameter, different flow rates, including 100, 200, 300, 400, and 500 μL/h, were tested.

### 2.5 RPA Combined with SE Chip for *Vibrio parahaemolyticus* Detection

In this research, Novec7500 containing surfactant was used as the continuous phase. Firstly, Novec7500 was injected into the chip to fill the entire reaction chamber and channels. Then, the RPA reagents were mixed and injected into the chip immediately using a syringe. The entire chip should be placed on ice to block the initiation of reaction. The chip containing the RPA system was immediately transferred to a heating plate set at 39°C for 30 minutes for isothermal amplification. Finally, the result was observed under a fluorescence microscope. The images were captured and the numbers of positive and negative droplets were counted by ImageJ. DNA molecules concentrations in the initial sample were calculated based on the Poisson distribution formula:

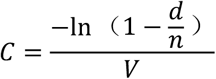

Where d is the number of positive droplets, n is the total number of droplets, and V is the volume of a single droplet. Quantitative detection of Vibrio parahaemolyticus was performed by using the SE chip combined with RPA and specificity and sensitivity were analyzed as described above.

### 2.6 Data Analysis

Statistical analysis were performed by using SPSS 20.0 software, conducting analysis of variance (ANOVA) and utilizing the Least Significant Difference (LSD) method for multiple group comparisons. Statistical significance is considered when P < 0.05.

## 3 Results

### 3.1 Specificity and Sensitivity Analysis of Traditioanl RPA for *Vibrio parahaemolyticus* Detection Traditional RPA assay was performed to verify the specificity of probe and primers

Amplification using primers and exo probe targeting *Vibrio parahaemolyticus* was performed on six pathogenic bacteria: *Vibrio parahaemolyticus* (Vp), *Proteus* (Pro), *Vibrio aerogenes* (Va), *Escherichia coli* (*E*.*coli*), *Vibrio alginolyticus* (Vg), and *Pseudomonas aeruginosa* (Pa) to validate the system’s specificity. The results show that only Vibrio parahaemolyticus exhibited significant amplification, with no significant amplification observed for the other bacteria, demonstrating good specificity (Figure 2a). The sensitivity of traditional RPA assay for *Vibrio parahaemolyticus* was also evaluated as described above, with various template concentrations. The DNA of target bacteria with an original concentration of 10^5^ CFU/μL was serially diluted to 10^4^, 10^3^, 10^2^, 10, and 1 CFU/μL. A negative control was included.

**Fig. 2.**
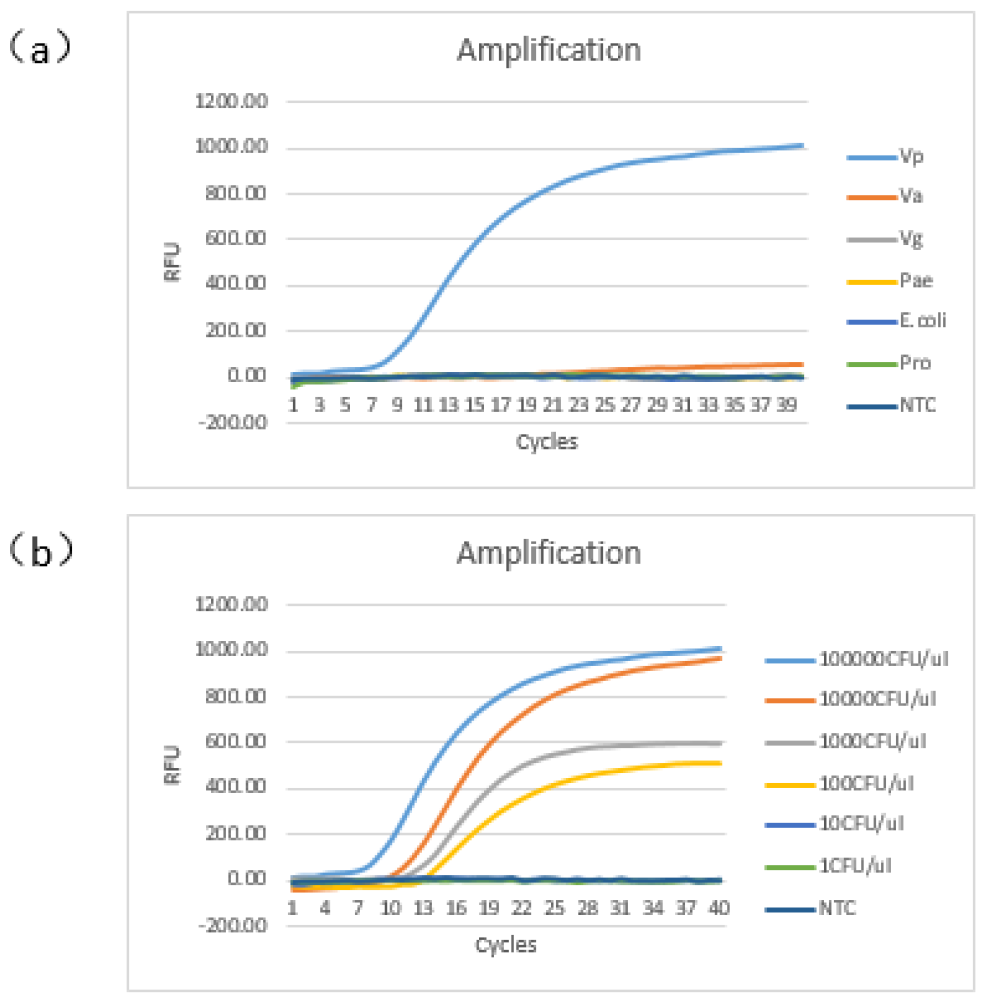
Reaction specificity and sensitivity of routine RPA

Real-time amplification results (figure 2b) indicated that conventional RPA could amplify the target gene when the initial template concentration ranged from 10^5^ to 10^2^ CFU/μL. However, amplification curves were not obtained at lower concentrations such as 10 and 1 CFU/μL. Therefore, the conventional RPA has a lowest detection limit of 10^2^ CFU/μL for *Vibrio parahaemolyticus*.

These findings suggest that the conventional RPA assay is specific to *Vibrio parahaemolyticus* and can detect the VP with sensitivity down to 10^2^ CFU/μL, making it a promising tool for rapid and accurate detection in various applications.

### 3.2 Characterization of Step Emulsification Chip

#### 3.2.1 Relationship between the channel height and droplet diameter

SE chips with different channel heights (10, 20,30,40, 50 μm, respectively) were fabricated and tested. Figure 3 illustrates the relationship between channel height and droplet diameter. The results revealed that as the channel height increases, the droplet diameter also increases accordingly. The results also demonstrated a proportional relationship between droplet diameter and channel height. A channel height of 40 μm was selected for subsequent experiments, and statistical analysis of 2000 droplets diameters at this height showed an average diameter of approximately 129 μm (Figure 4).

**Fig. 3.**
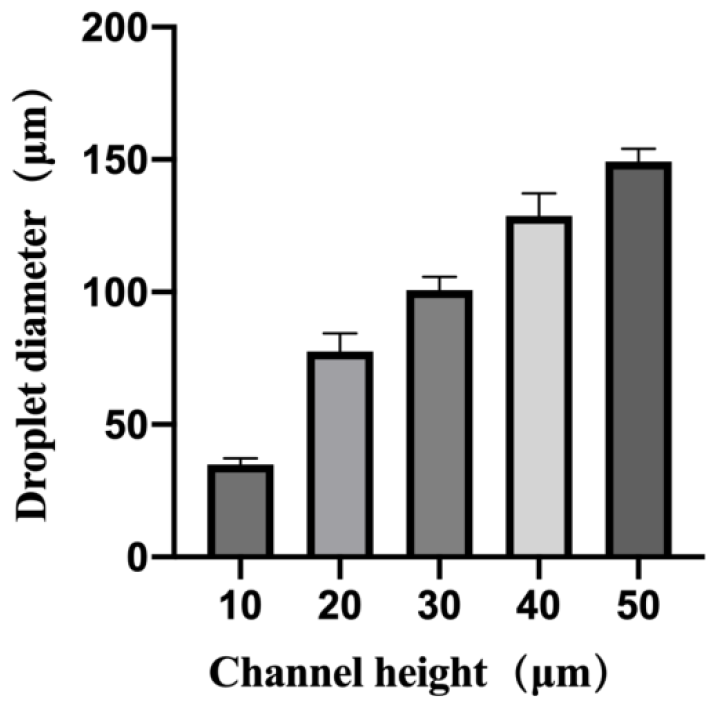
The effect of chip channel height on droplet size

**Fig. 4.**
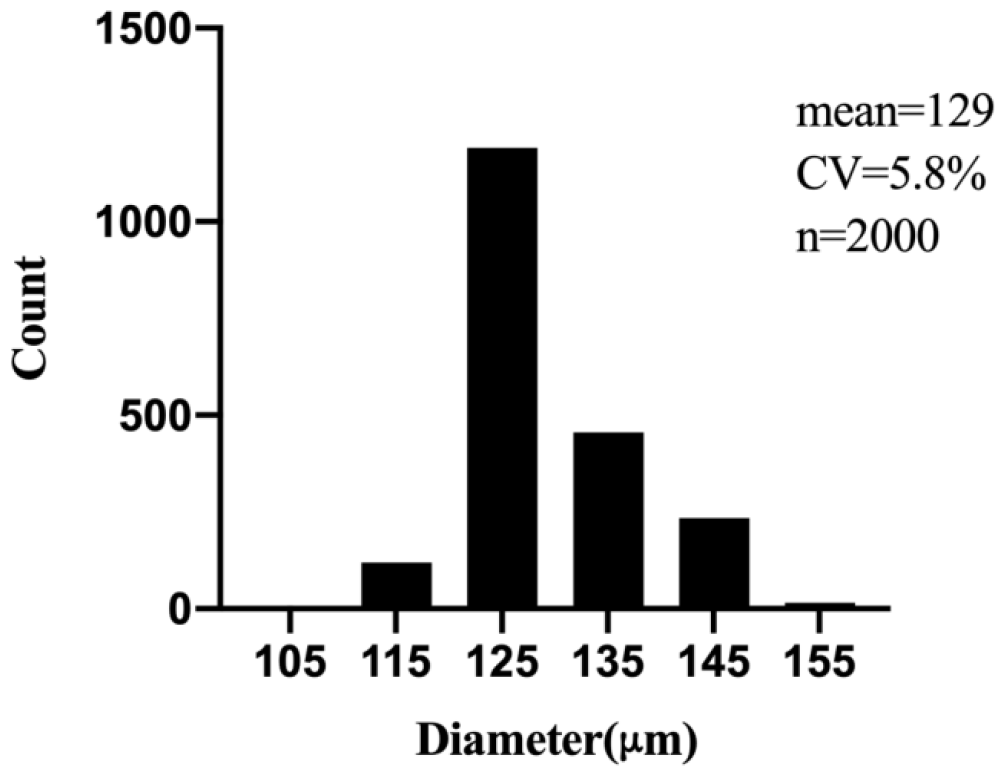
Diameter distribution of droplets in 40μm-SE chip

#### 3.2.2 Flow Rate-Droplet Diameter Relationship

As shown in Figure 5, when the channel height is 40 μm, the effect of dispersed phase flow rates (100, 200, 300, 400, and 500 μl/h) on droplet diameter were tested. Results indicated that droplet diameters ranged from 128 to 132 μm under different flow rates. Combined with results in section 4.2.1, it was demonstrated that droplet diameter primarily depends on the chip’s channel height, while flow rates of dispersed phase having a minor impact.

**Fig. 5.**
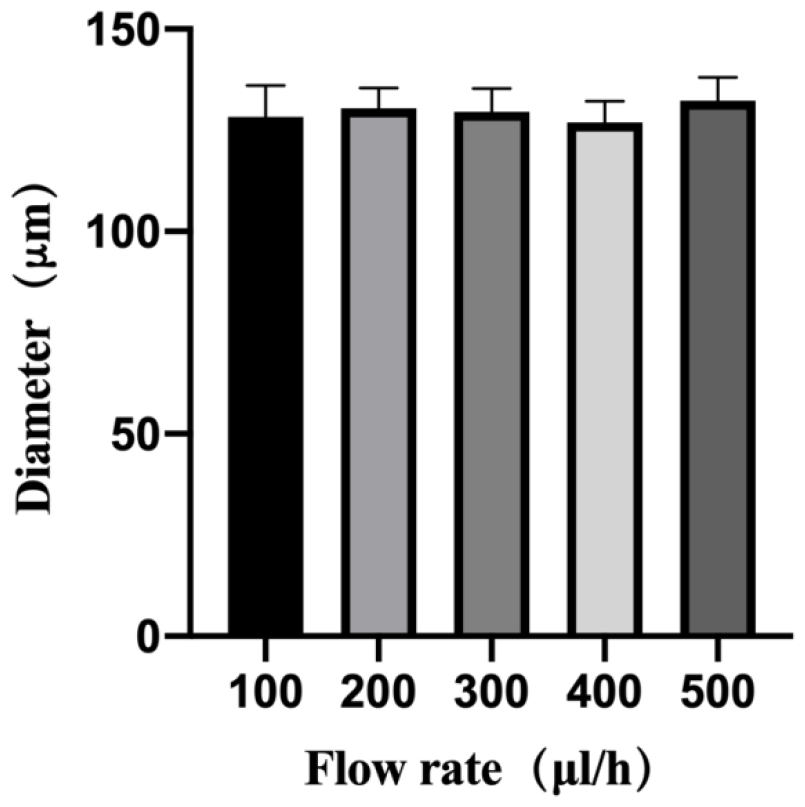
The influence of flow rate on the droplets diameter

### 3.3 Detection of *Vibrio parahaemolyticus* Using RPA Combined with SE Chip

#### 3.3.1 Changes in Droplet Fluorescence Intensity with Incubation Time

During the reaction, fluorescence images of droplets were captured every 5 minutes using a fluorescence microscope. Figure6 show microscopic images of droplets at incubation times of 0, 10, 20, 30, and 40 minutes, with a scale of 200 μm. It was observed that the fluorescence of positive droplets gradually increased with increasing incubation time.

**Fig. 6.**
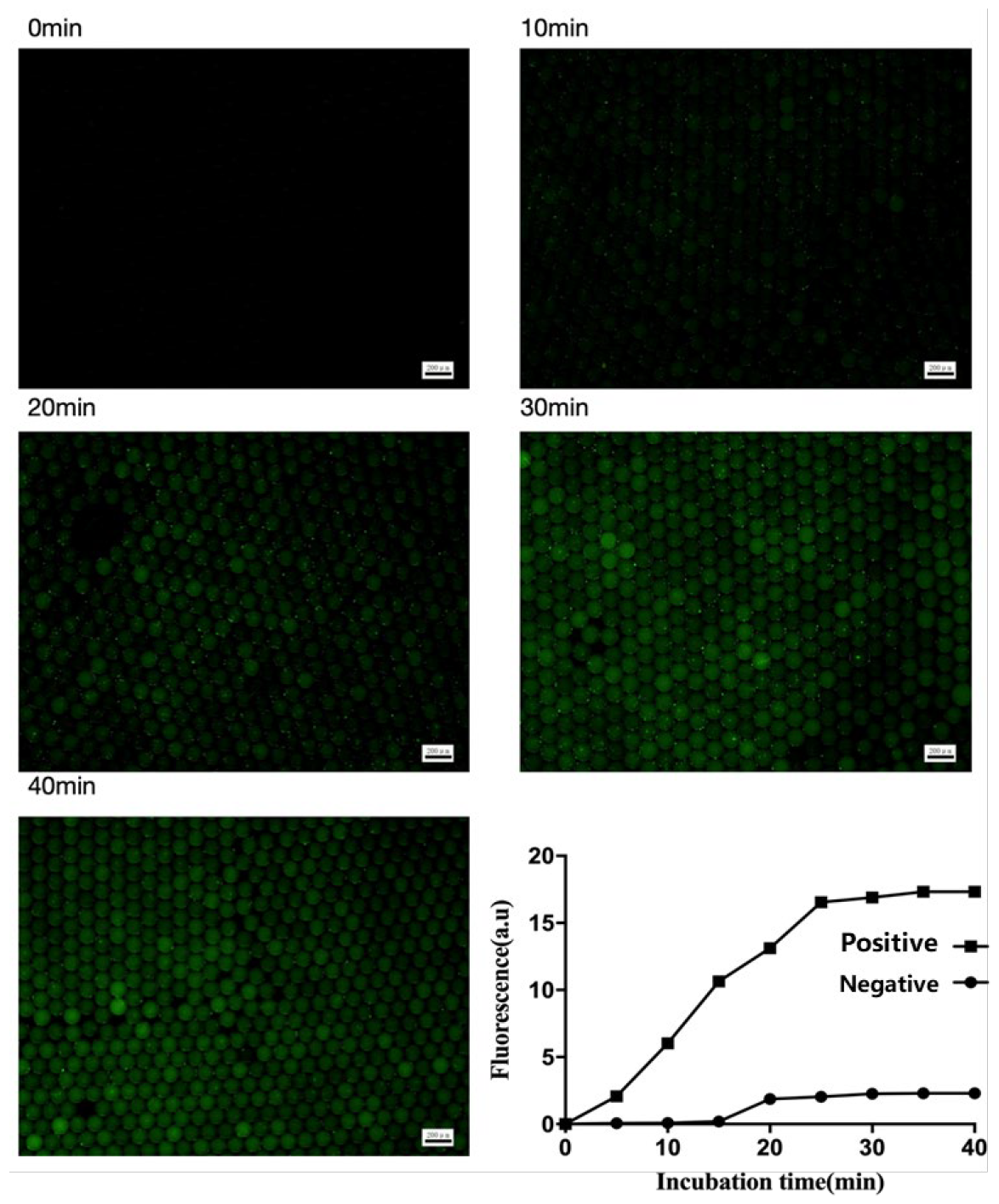
Change in the fluorescence intensity of the droplets during the incubation

Analysis of the fluorescence intensity of positive droplets using Image J (Figure 7) revealed that compared to negative droplets, the real-time fluorescence intensity of positive droplets increased. When the amplification time reached 25 minutes, the fluorescence intensity approached a plateau. To optimize reaction time, an incubation time of 25 minutes was selected for subsequent experiments.

**Figure 7.**
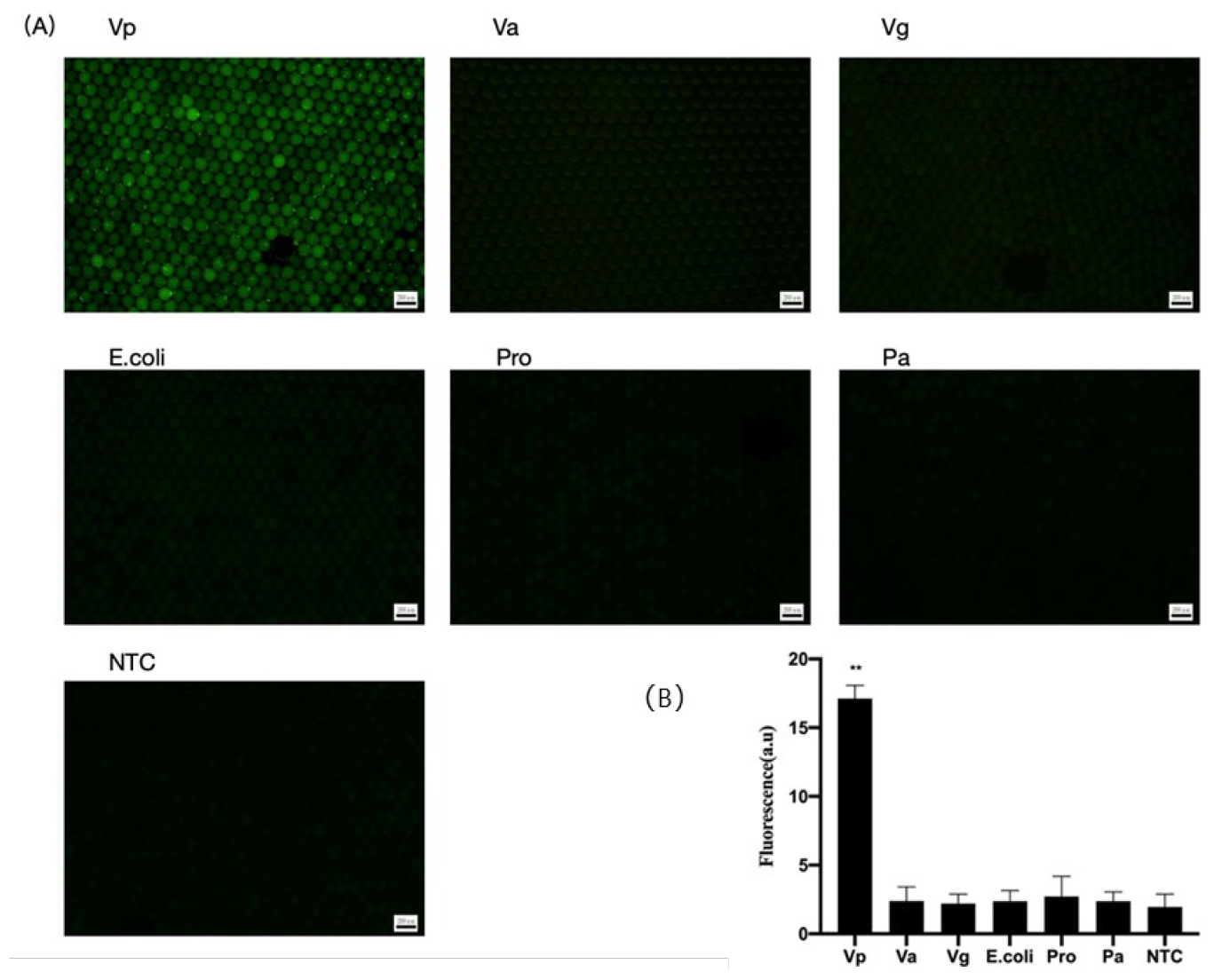
Specificity tests of the on-chip RPA

#### 3.3.2 Specificity Analysis

The same amplification system was used to conduct droplet RPA reactions with six different target bacteria on the chip. It was found that *Vibrio parahaemolyticus* produced positive droplets, while other bacteria did not yield positive droplets, demonstrating successful amplification of Vp with good specificity (Fig.7).The fluorescence intensities were also analyzed (Fig.7B). The system successfully amplified *Vibrio parahaemolyticus*, distinguishing it from other target bacteria, with **P<0.01 vs the control group, indicating statistical significance.

#### 3.3.3 Sensitivity Analysis

Absolute quantification of different concentrations (10^4^, 10^3^, 10^2^, 10^1^ CFU/μL) of *Vibrio parahaemolyticus* were performed using RPA combined with the SE chip. The results showed that, as the DNA concentration decreases, the number of positive droplets decreases. Calculations based on the Poisson distribution formula for the number of positive droplets relative to negative droplets yielded initial sample concentrations consistent with expected values, with an R^2^ value of 0.9499 (Figure 8). This sensitivity analysis demonstrates that RPA combined with the SE chip allows for accurate and quantitative detection of *Vibrio parahaemolyticus* across a wide range of concentrations, providing reliable results for practical applications requiring precise pathogen quantification.

**Figure 8.**
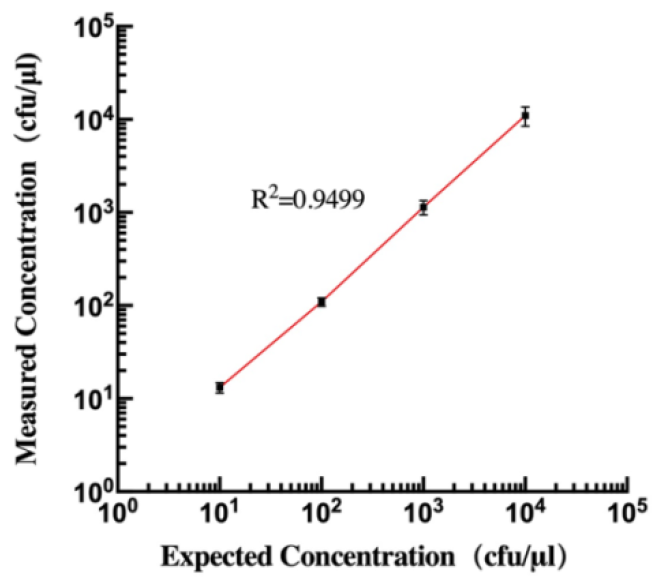
sensitivity of SE RPA

## 4. Discussion

*Vibrio parahaemolyticus*, a Gram-negative halophilic marine pathogen, mainly inhabits marine and river mouth. Since its discovery in 1950, *Vibrio parahaemolyticus* has been recognized as the major cause of seafood-borne diseases.^17-19^ Despite the availability of various techniques for *Vibrio parahaemolyticus* detection, there is still a demand for a simple, user-friendly, and highly sensitive quantitative testing platform. ^20-24^

In this research, microfluidic was used as the detection platform, employing isothermal amplification RPA technology to amplify *Vibrio parahaemolyticus*, enabling rapid detection process. A microfluidic SE chip is used to produce monodisperse droplets. Isothermal amplification technology is employed to incubate the droplets inside the chip, followed by observation of fluorescent results. Compared to traditional methods such as T-channel and flow-focusing techniques, the SE structure offers advantages in terms of reliability and ease of fabrication. The size of the droplets is mainly determined by the geometric structure of the chip. Once the structure is established, the droplet size produced by the chip is also fixed. Therefore, the SE chip can simultaneously utilize parallel large-scale nozzle arrays to achieve high-throughput droplet production and have been widely used many fields.^25^ In this research, droplets are used to divide the reaction system into tens of thousands of oil-encapsulated water droplets, with each droplet representing a conventional RPA reaction system. By using an *exo* probe in the system, negative and positive droplets can be distinguished, allowing for absolute quantification of the initial sample concentration of *Vibrio parahaemolyticus*. This method features characteristics such as no contamination, low sample consumption, high digitization, high throughput, and high stability. Notably, we found that the detection limit in digital-RPA (10 CFU/μL) is 10 times lower than that in traditional tube-based RPA (100 CFU/μL), which may be due to reducing background noise and increasing the signal-to-noise ratio in digital-RPA.^26^

This research provided a step emulsion microfluidic chip. Combined with RPA technology, the *Vibrio parahaemolyticus* was detected. This new method provided many advantages such as labor-saving, easy to operate and improved sensitivity, among others.

